# Genome Assembly of the Endangered Patagonian Deer *Hippocamelus bisulcus* (huemul): The First Nuclear Genome for the Genus *Hippocamelus*

**DOI:** 10.64898/2026.09.04.747633

**Authors:** María Julia Ousset, Jo Anne M. Smith-Flueck, Werner T. Flueck, Valeria Pelufo, Eduardo Aisen, Andrés Venturino

## Abstract

The huemul (*Hippocamelus bisulcus*) is an endangered cervid endemic to the Andean–Patagonian region of South America, where it persists in small, fragmented populations. The lack of a reference genome has limited genomic approaches to huemul conservation and evolutionary research. Despite moderate theoretical coverage (∼22.8×), Oxford Nanopore long reads yielded the first highly contiguous and nearly complete nuclear genome assembly for *H. bisulcus*. The 2.50-Gb assembly achieved a contig N50 of 8.75 Mb, 99.0% genome-mode BUSCO completeness, an estimated k-mer completeness of 95.63%, and an ONT k-mer-based QV estimate of 48.11. Reference-guided scaffolding against the white-tailed deer (*Odocoileus virginianus*) genome organized 94% of the assembly into 36 chromosome-scale pseudomolecules (34 autosomes, X, and Y; scaffold N50 = 68.56 Mb). Repeat annotation identified 38.09% of the assembly as repetitive, dominated by LINEs, consistent with other cervid genomes. Coordinate-based annotation transfer with LiftOn identified 20,042 protein-coding genes, with 95.9% protein-mode BUSCO completeness in the representative predicted proteome. We also assembled a complete circular mitochondrial genome of 16,405 bp containing the expected 37-gene complement in the conserved vertebrate arrangement. Nuclear and mitochondrial phylogenies placed *H. bisulcus* within Odocoileini (Capreolinae), while the mitochondrial analysis recovered *H. bisulcus* and the taruka, *H. antisensis*, as a maximally supported sister pair. Comparative analysis across nine Cervidae proteomes assigned 98.8% of the representative *H. bisulcus* proteins to orthogroups shared with at least one other species, indicating broad recovery of the conserved cervid protein repertoire. This study provides the first nuclear genome for the South American genus *Hippocamelus* and the first nuclear and mitochondrial genomic resources for *H. bisulcus*, establishing a foundational framework for population genomics, conservation management, and evolutionary studies of this emblematic Patagonian deer.

## Introduction

The huemul (*Hippocamelus bisulcus*; Fig. 1, right) is an endangered cervid endemic to the Southern Cone of South America. During the Holocene, the species occupied a much broader distribution, extending nearly 2,800 km from approximately 30°S to 55°S in Argentina and from the Patagonian Andean range eastward to the Atlantic coast (Smith-Flueck et al., 2025; Fig. 1, left). Post-Columbian anthropogenic pressures, including overhunting, land conversion, habitat fragmentation, and the introduction of domestic livestock, drove a severe contraction of its historical range (Flueck et al., 2022, 2023; Huemul Task Force, 2012; Zuliani et al., 2023). Today, approximately 1,500 individuals remain in more than 100 fragmented subpopulations, largely restricted to the Andes of Chile and Argentina. Due to loss of migratory traditions, most extant populations now persist year-round in mountain refugia that historically represented seasonal summer range. The situation is further exacerbated by trace mineral deficiencies in soils of upper elevations that severely affect the general population’s health, ultimately restricting population growth (Flueck et al., 2022; Smith-Flueck et al., 2025).

**Figure 1.**
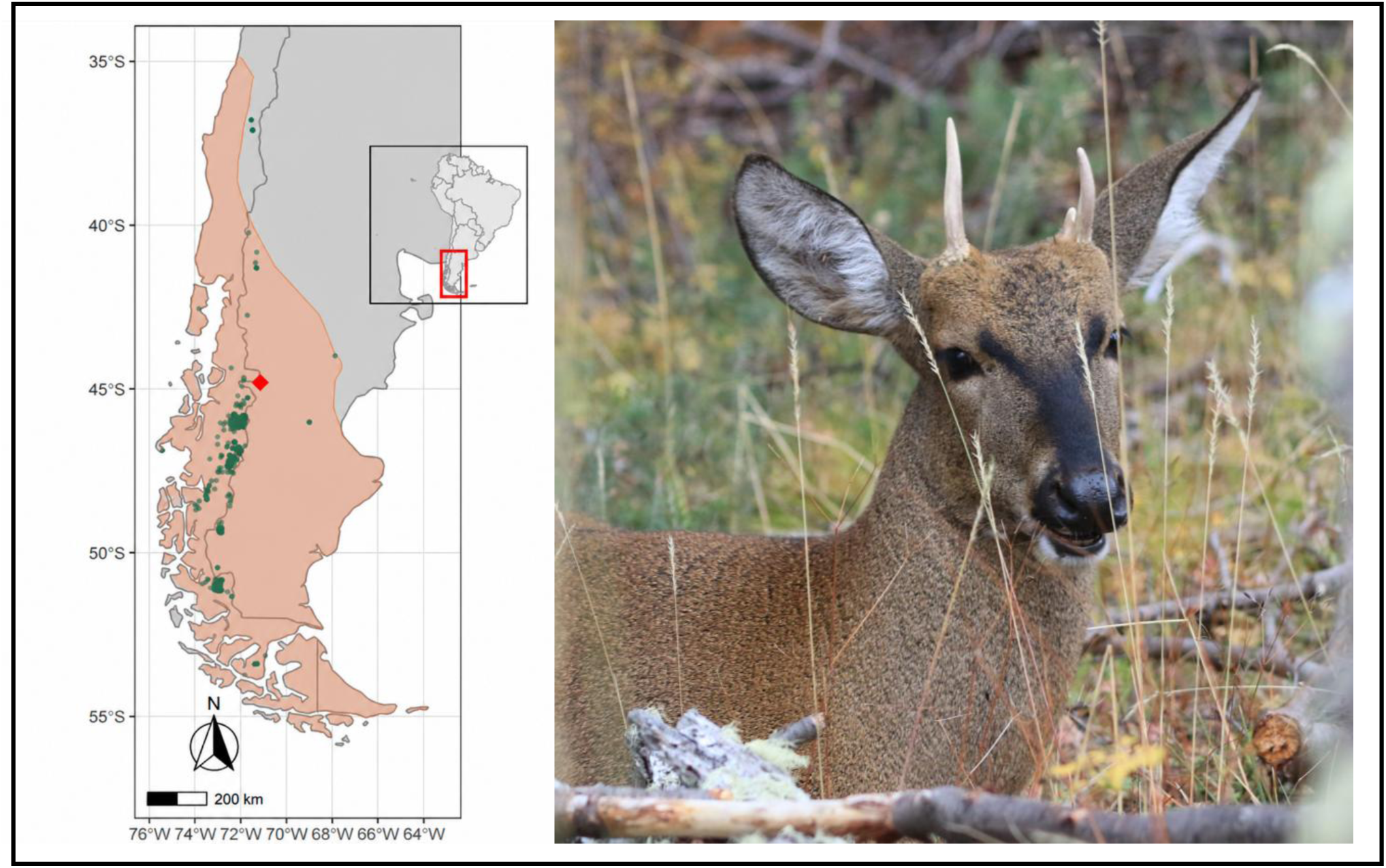
Sampled individual and geographic context of the huemul (*Hippocamelus bisulcus*). Left: historical distribution of *H. bisulcus* in southern South America (shaded area). Green dots indicate contemporary occurrence records, and the red diamond marks the sampling locality of the two individuals included in this study. The inset shows the location of the mapped area within South America. Distribution adapted from Smith-Flueck et al. (2025). Right: Shehuen, a yearling male huemul that provided the principal genomic DNA sample for the nuclear genome assembly and the complete mitochondrial genome reported in this study. Photograph by Jo Anne Smith-Flueck.

The management of small, fragmented populations increasingly relies on conservation genomic approaches to assess inbreeding, effective population size, connectivity, and adaptive potential (Hohenlohe et al., 2021). The availability of a whole-genome reference substantially expands the range of questions that can be addressed, enabling genome-wide analyses of inbreeding, demographic history, population connectivity, local adaptation, and evolutionary relationships (Brandies et al., 2019; Supple & Shapiro, 2018; Theißinger et al., 2023). In cervids, genomic resources have expanded considerably in recent years, with reference genomes now available for northern Odocoileini such as the white-tailed deer (*Odocoileus virginianus*) (London et al., 2022) and mule deer (*O. hemionus*) (Lamb et al., 2021). However, genomic resources remain sparse and unevenly distributed across Cervidae (Zhong et al., 2026), and no nuclear reference genome has previously been available for *Hippocamelus*. For *H. bisulcus* specifically, genetic studies have to date relied primarily on mitochondrial markers and microsatellites (Corti et al., 2011; Smith-Flueck et al., 2025), approaches that have long provided valuable insights into population history, phylogeography, and evolutionary relationships in non-model species (Galtier et al., 2009).

Oxford Nanopore Technologies (ONT) long-read sequencing offers particular advantages for developing genomic resources in non-model and threatened species. Long reads facilitate *de novo* genome assembly and improve contiguity by spanning repetitive and structurally complex regions, while the portability and relatively low infrastructure requirements of nanopore sequencing allow genomic data to be generated closer to the source of biological samples (Hauff et al., 2025; Wang et al., 2021). This combination can reduce dependence on centralized sequencing facilities and, in some cases, the need to transport or export biological material across national borders, an important consideration when working with protected species (Hauff et al., 2025; Pozo et al., 2024). These advantages have already been demonstrated in threatened non-model species from biodiversity-rich regions, including primates in Ecuador (Pozo et al., 2024) and Madagascar (Hauff et al., 2025). For the huemul, whose populations occur in remote Patagonian environments and whose biological material is subjected to logistical and regulatory constraints, the possibility of generating long-read genomic data locally is particularly valuable.

In this Genetics Note, we present the first reference genome assembly for *H. bisulcus*, a highly complete 2.50-Gb resource organized into chromosome-scale pseudomolecules through reference-guided scaffolding. We describe the nuclear genome assembly and structural annotation, together with the complete mitochondrial genome and comparative phylogenomic analyses within Cervidae. This study establishes the first nuclear genomic resource for the genus *Hippocamelus*, while also providing the first nuclear and mitochondrial genomic resources for *H. bisulcus*, and lays the foundation for future evolutionary, population, and conservation genomics studies and genomics-informed management of the huemul.

## Materials and Methods

All bioinformatic analyses were performed locally, except for specific steps carried out on the Galaxy platform (The Galaxy Community, 2024) on the public usegalaxy.eu server, which are indicated explicitly where relevant.

### DNA Extraction

Blood samples were obtained from two male *Hippocamelus bisulcus* individuals during routine veterinary and conservation-management procedures at the Shoonem Breeding and Rehabilitation Center, Chubut, Argentina. One of the males, Shehuen, was a juvenile (yearling) and the first huemul born in captivity in Argentina at the Shoonem Center (Figure 1). The second male, Coirón, was a wild-born young adult from the Lake La Plata subpopulation and was brought to the Center in 2023. High-molecular-weight genomic DNA was isolated from both blood samples using a CTAB-based protocol adapted from Murray and Thompson (1980). The DNA was subsequently purified using the Monarch® Spin gDNA Extraction Kit (New England Biolabs, Ipswich, MA, USA; catalog no. T3010), following the manufacturer’s instructions for genomic DNA cleanup to ensure the purity and integrity required for long-read sequencing.

### Nanopore Sequencing and Basecalling

A total of five genomic libraries were prepared using the Oxford Nanopore Technologies (ONT) Ligation Sequencing Kit V14 (SQK-LSK114), four using DNA from Shehuen and one using DNA from Coirón. Sequencing was performed on an ONT MinION Mk1B platform using eight R10.4.1 flow cells, following the manufacturer’s instructions. Data acquisition was managed with MinKNOW software. Raw signal data were basecalled using Dorado within the epi2me-labs/wf-basecalling pipeline v1.5.9 (EPI2ME Labs, 2026; Ewels et al., 2020) in super-accuracy mode with the R10.4.1 model dna_r10.4.1_e8.2_400bps_sup@v5.2.0. Adapter trimming was performed during basecalling using Dorado’s built-in trimming functionality (Oxford Nanopore Technologies, 2026a), and reads were filtered to retain sequences with a minimum quality score of 10. The basecalled and quality-filtered reads from Shehuen and Coirón were deposited separately in the NCBI Sequence Read Archive (SRA) under BioProject PRJNA1509444, with accession numbers SRR40078882 and SRR40078881, respectively. The corresponding BioSample accessions are SAMN62264853 for Shehuen and SAMN62264854 for Coirón.

### Mitochondrial genome

#### Mitochondrial assembly and annotation

To assemble the mitochondrial genome of *H. bisulcus*, long reads from Shehuen were first size-selected with Filtlong v0.3.1 (Wick, 2017), retaining only reads between 5,000 and 16,500 bp to enrich mitochondrial-length molecules. The filtered reads were then mapped against the *Hippocamelus antisensis* complete mitochondrial genome (NC_020711.1) to enrich for reads of mitochondrial origin, using minimap2 v2.31 (Li, 2018) with the map-ont preset, with secondary alignments disabled, a minimum secondary-to-primary score ratio of 0.9, and a minimum peak DP alignment score of 100 to reduce co-mapping of nuclear mitochondrial DNA segments (NUMTs). Reads mapping to the reference were extracted and converted to FASTQ with SAMtools v1.23.1 (Danecek et al., 2021) for *de novo* assembly. The mitogenome was assembled from the enriched reads using PMAT2 v2.1.5 (Han et al., 2025) in autoMito mode, specifying ONT input, mitochondrial genome assembly, and animal taxonomy. Within the PMAT2 workflow, reads were error-corrected with Canu v2.2 (Koren et al., 2017) prior to assembly, and an expected mitochondrial genome size of 16,400 bp was provided to bypass genome-size estimation from the mitochondrial-enriched read set. Structural annotation was then performed with MITOS2 v2.1.0 (Donath et al., 2019) run on the Galaxy platform, using the vertebrate mitochondrial genetic code (NCBI translation table 2). The complete mitochondrial sequence was submitted to GenBank as an organellar sequence within WGS project JCCBWI000000000 (BioProject PRJNA1509444).

### Mitochondrial phylogenetic analysis

For the mitochondrial phylogeny, a total of 16 complete mitochondrial genomes were analyzed, comprising representatives of multiple cervid genera together with a bovid as an outgroup. Four mitochondrial replicons were retrieved from chromosome-level genome assemblies used in the proteome-based comparative analysis conducted in this study, where an annotated mitochondrial sequence was available: *Cervus canadensis* (CM033226.1), *Cervus elaphus* (OU343111.2), *Dama dama* (CM065635.2), and *Muntiacus reevesi* (OZ005647.2). The remaining cervid sequences were selected as one representative per genus from the NCBI Nucleotide RefSeq database: *Hippocamelus antisensis* (NC_020711.1), *Ozotoceros bezoarticus* (NC_020766.1), *Blastocerus dichotomus* (NC_020682.1), *Pudu puda* (NC_020740.1), *Mazama americana* (NC_020719.1), *Odocoileus virginianus* (NC_015247.1), *Rangifer tarandus* (NC_007703.1), *Alces alces* (NC_020677.1), *Capreolus capreolus* (NC_020684.1), and *Muntiacus muntjak* (NC_004563.1). The newly assembled *H. bisulcus* mitogenome was included, and *Bos taurus* (NC_006853.1) was used as the outgroup. All mitochondrial sequences were checked for orientation and standardized to the same genomic direction. The newly assembled *H. bisulcus* mitogenome was additionally rotated so that position 1 corresponded to the conventional mitochondrial genome starting point used in the reference sequences prior to alignment. Multiple sequence alignment was performed with the MAFFT v7 online server (Katoh et al., 2019), with automatic strategy selection and direction adjustment enabled. Maximum-likelihood phylogenetic inference was carried out with IQ-TREE v3.1.2 (Wong et al., 2026). The best-fit substitution model, TIM2+F+G4, was selected by ModelFinder according to the Bayesian information criterion (BIC), and branch support was estimated from 1,000 ultrafast bootstrap replicates using UFBoot2 (Hoang et al., 2018; Kalyaanamoorthy et al., 2017). The resulting tree was visualized using a custom Python script with the Biopython v1.84 Phylo module (Cock et al., 2009) and Matplotlib v3.9 (Hunter, 2007).

### Nuclear genome assembly

Exploratory assembly tests were carried out on the Galaxy platform to identify the optimal assembly strategy. These included assembly from a single individual (Shehuen, 48 Gb of ONT reads) versus combined data from both individuals (Shehuen + Coirón, 56 Gb of ONT reads), alternative read-filtering strategies, variation of the minimum read-overlap parameter in Flye, and *de novo* assembly with Flye and Hifiasm, including haplotype-aware and non-haplotype-aware configurations. Candidate assemblies were compared using QUAST and BUSCO on Galaxy. These exploratory comparisons were used solely to optimize the workflow and identify the best-performing assembly strategy. Based on these comparisons, the assembly generated with Flye v2.9.6 (Kolmogorov et al., 2019) from the combined long-read dataset was selected for all subsequent processing and evaluation and is the assembly reported in this study. Flye was run in --nano-hq mode, with a minimum read overlap of 1,000 bp and a single polishing iteration. The underlying read datasets are available separately in the NCBI Sequence Read Archive under BioProject PRJNA1509444, allowing both the single-individual and combined-data assembly strategies to be reproduced.

All subsequent assembly processing and characterization were performed locally. Assembly statistics and contiguity were evaluated with QUAST v5.3.0 (Gurevich et al., 2013), and gene-space completeness was assessed with BUSCO v6.1.0 (Manni et al., 2021) against the cetartiodactyla_odb10 lineage dataset.

### Post-assembly processing

#### Haplotypic Duplication Removal

Haplotypic duplications and redundant contig overlaps were removed using purge_dups v1.2.5 (Guan et al., 2020), following the standard pipeline. Long reads from both individuals were aligned to the assembly with minimap2 v2.31 (Li, 2018) using the Oxford Nanopore read-mapping preset. Base-level read-depth profiles were generated with pbcstat, followed by automatic cutoff estimation with calcuts, which yielded an upper coverage cutoff of 66×. The assembly was split and self-aligned with minimap2 in assembly-to-assembly mode to detect overlapping and duplicated contigs. The resulting self-alignment file was analyzed with purge_dups together with the coverage data and estimated cutoffs, using the two-round chaining option. Finally, get_seqs was run with the end-specific option, yielding the purged primary assembly.

### Polishing

The purged primary assembly was polished with a single round of Medaka v2.2.2 (Oxford Nanopore Technologies, 2026b) using the consensus pipeline and the r1041_e82_400bps_sup_v5.0.0 model. A k-mer database was generated from the combined quality-filtered ONT reads of both individuals using Meryl v1.4.1 with a k-mer size of 21 (Rhie et al., 2020). This database was used with Merqury v1.3 under default settings to compare the raw Flye assembly with the Medaka-polished assembly and to estimate changes in consensus quality and k-mer completeness after polishing (Rhie et al., 2020).

### Contamination removal

The polished assembly was screened for contamination in Galaxy using NCBI FCS-GX with the fcs-2023-01-24 database in screen mode (Astashyn et al., 2024). Contigs classified as contaminants in the FCS-GX action report were subsequently inspected and removed locally by filtering their sequence identifiers with SeqKit v2.13.0 (Shen et al., 2016). The resulting assembly was then screened for mitochondrial homology by aligning it against a doubled copy of the previously assembled *H. bisulcus* mitochondrial genome, used to account for mitochondrial genome circularity and avoid edge effects at the linearized sequence boundaries, using BLASTn from the BLAST+ v2.17.0 suite, in megablast mode, with an E-value threshold of 0.001 and DUST low-complexity filtering enabled (Camacho et al., 2009). Subsequently, the assembly was filtered with SeqKit v2.13.0 to retain only contigs of at least 20,000 bp. As a final pre-scaffolding curation step, the length-filtered assembly was screened for residual adaptor and vector contamination using NCBI FCS-adaptor v0.5.0 on the Galaxy platform in eukaryotic mode. The resulting adaptor action report was applied using the NCBI FCS-GX v0.5.5 clean-genome procedure, with a minimum retained sequence length of 200 bp, to trim or exclude regions flagged by FCS-adaptor and generate the cleaned assembly (Astashyn et al., 2024).

### Scaffolding

The final curated assembly was scaffolded against the *Odocoileus virginianus* reference genome (Ovbor_1.2, GCF_023699985.2) using RagTag v2.1.0 in scaffold mode (Alonge et al., 2022). Minimap2 v2.31 (Li, 2018) was used as the alignment engine with the asm5 assembly-alignment preset and a minimum minimizer-occurrence floor of 100. Unique alignments shorter than 1,000 bp were excluded, and a minimum mapping quality of 10 was required. Default confidence thresholds were used for grouping, placement, and orientation. Gaps between adjacent contigs were assigned the default size of 100 bp. Unplaced contigs were initially concatenated by RagTag into a single chr0 sequence. Prior to GenBank submission, the composite chr0 sequence was decomposed into its constituent unplaced contigs using the component coordinates and orientations recorded in the RagTag AGP file. Each component was extracted from chr0, reverse-complemented when required by its AGP orientation, and retained as an independent sequence in the final assembly. The nuclear genome assembly has been deposited at DDBJ/ENA/GenBank under the accession JCCBWI000000000. The version described in this paper is JCCBWI010000000.

### Repeat Annotation

Repeat annotation was performed using the Galaxy platform. *De novo* repeat families were identified with RepeatModeler v2.0.5 using default parameters (Flynn et al., 2020), and the resulting custom repeat library was subsequently used to soft-mask the final scaffolded assembly deposited in GenBank with RepeatMasker v4.1.5 (Smit et al., 2013). RepeatMasker was run with a cutoff score of 225, maximum sequence fragment length of 40 kb, exclusion of N-runs from repeat statistics, and soft-masking enabled.

### Structural Annotation

Gene models from the *Odocoileus virginianus* reference genome (Ovbor_1.2, GCF_023699985.2; NCBI RefSeq annotation release 2024-12) were transferred to the unmasked scaffolded *Hippocamelus bisulcus* assembly using LiftOn v1.0.9 (Chao et al., 2025). Prior to annotation transfer, the mitochondrial sequence NC_015247.1 and its associated annotations were removed from the *O. virginianus* reference FASTA and GFF3 files to prevent spurious transfer of mitochondrial genes onto nuclear mitochondrial DNA segments (NUMTs) in the *H. bisulcus* assembly. LiftOn was then run with the extra-copy option enabled and a minimum sequence-identity threshold of 0.95. In parallel, an independent *ab initio* annotation was generated in Galaxy using BRAKER3 v3.0.8 (Gabriel et al., 2024) with *O. virginianus* protein sequences combined with publicly available RNA-seq data from two *O. virginianus* tissues retrieved from NCBI SRA: muscle (SRR4256031) and liver (SRR4256028), both from the White-tailed Deer Genome Project. Protein sequences for the BRAKER3 annotation were obtained from the native protein output generated by BRAKER3, whereas LiftOn protein sequences were extracted from the transferred annotation using AGAT v1.7.0 (Dainat et al., 2026). For both annotations, the longest protein isoform per gene was retained to generate representative proteomes and avoid duplication caused by alternative transcripts. Annotation completeness was assessed on these representative protein sets using BUSCO v6.1.0 in protein mode with the cetartiodactyla_odb10 lineage dataset (Manni et al., 2021). General annotation statistics for both LiftOn and BRAKER3 were calculated from their respective complete GFF3 annotations using AGAT v1.7.0. The two annotation strategies were compared based on representative-proteome BUSCO completeness, and the LiftOn-derived annotation was retained as the primary structural annotation.

### Comparative Genomics and Phylogenetic Analysis

For comparative genomic analysis, Cervidae taxa with publicly available annotated proteomes were selected from NCBI, together with *Bos taurus* for phylogenetic rooting. The dataset comprised the newly annotated *Hippocamelus bisulcus* proteome and *Cervus canadensis* (GCF_019320065.1), *Cervus elaphus* (GCF_910594005.1), *Cervus hanglu yarkandensis* (GCA_010411085.1), *Dama dama* (GCF_033118175.1), *Muntiacus muntjak* (GCA_008782695.1), *Muntiacus reevesi* (GCF_963930625.1), *Odocoileus virginianus* (GCF_023699985.2), *Rangifer tarandus platyrhynchus* (GCA_949782905.1), and *Bos taurus* (GCF_002263795.3). To reduce isoform redundancy, each proteome was standardized by retaining the longest protein isoform per gene before OrthoFinder analysis. To identify single-copy orthologues and generate a concatenated species-tree alignment, OrthoFinder v3.1.5 was run in multiple-sequence-alignment mode using DIAMOND for protein similarity searches, FAMSA for multiple-sequence alignment generation, and FastTree for gene-tree inference (Buchfink et al., 2021; Deorowicz et al., 2016; Emms & Kelly, 2019; Price et al., 2010). The resulting concatenated alignment of single-copy orthologues was subsequently used as input for phylogenetic inference with IQ-TREE v3.1.2. The best-fit substitution model, Q.MAMMAL+F+I+R7, was selected by ModelFinder according to the Bayesian information criterion (BIC), and branch support was estimated from 1,000 ultrafast bootstrap replicates using UFBoot2 (Hoang et al., 2018; Kalyaanamoorthy et al., 2017; Wong et al., 2026). The IQ-TREE consensus tree was relabeled using the OrthoFinder species-ID mapping. For visualization, *Bos taurus* was designated as the outgroup, and the tree was rooted using the root_with_outgroup function from Biopython v1.84 (Cock et al., 2009). The rooted tree was subsequently visualized with Matplotlib v3.9 (Hunter, 2007).

The complete genome assembly, curation, and annotation workflow, together with summary statistics at each processing stage, is shown in Figure 2.

**Figure 2.**
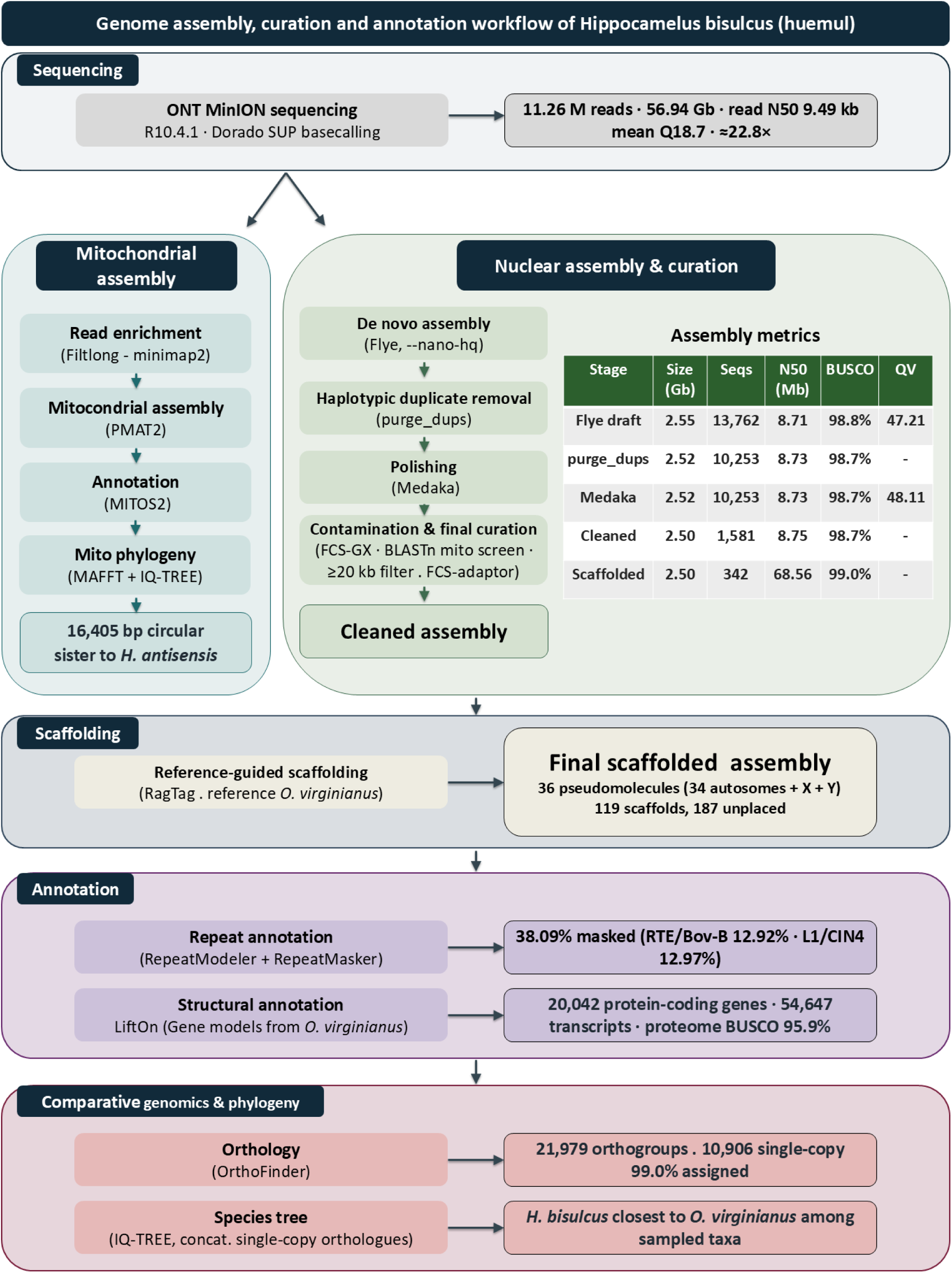
Overview of the *H. bisulcus* genome assembly, curation, and annotation workflow. Each panel shows the main tool(s) at each stage, with key outputs alongside; the table summarizes assembly metrics from the Flye draft to the final scaffolded assembly.

## Results

### Nanopore Sequencing and Basecalling

Eight MinION R10.4.1 flow-cell runs generated approximately 632 GB of raw signal data in POD5 format, comprising approximately 523 GB from Shehuen and 108 GB from Coirón. Following super-accuracy basecalling and quality filtering (Q ≥ 10), a total of 11.26 million reads representing 56.94 Gb of sequence data were retained, with a read N50 of 9.49 kb and a mean read quality of Q18.7. Shehuen contributed 10.16 million reads (48.07 Gb), with a read N50 of 8.94 kb, whereas Coirón contributed 1.10 million reads (8.86 Gb), with a read N50 of 13.22 kb. The longest reads obtained were 602.95 kb for Shehuen and 303.04 kb for Coirón. These reads constituted the input dataset for the subsequent mitochondrial and nuclear genome analyses. The combined retained read set corresponded to an estimated theoretical nuclear genome coverage of approximately 22.8×, based on the final 2.50-Gb assembly size.

### Mitochondrial genome

Pre-assembly read enrichment against the *H. antisensis* reference mitochondrial genome (NC_020711.1), combined with stringent mapping parameters to minimize co-enrichment of nuclear mitochondrial DNA segments (NUMTs), yielded 1,297 reads for *de novo* mitochondrial genome assembly. PMAT2 recovered a single circular mitochondrial molecule of 16,405 bp, with an estimated assembly depth of approximately 40.5× based on the mitochondria-enriched read set. Structural annotation with MITOS2 identified the complete set of 37 genes expected for a vertebrate mitochondrial genome (Boore, 1999), comprising 13 protein-coding genes, 22 transfer RNA genes, and two ribosomal RNA genes, with no missing genes detected. Three gene overlaps were identified: *atp8/atp6* (40 bp), *nad4l/nad4* (7 bp), and *trnS2/cox1* (3 bp). A BLASTn search of the assembled mitogenome against publicly available cervid mitochondrial genomes identified the *H. antisensis* mitogenome (NC_020711.1) as the closest match, with 96.23% nucleotide identity and 100% query coverage. This phylogenetic affinity was further supported by maximum-likelihood analysis, which recovered *H. bisulcus* and *H. antisensis* as sister taxa with maximal support (ultrafast bootstrap = 100). Both species were nested within a South American clade of Odocoileini (ultrafast bootstrap = 84) that also included *Blastocerus dichotomus* and *Ozotoceros bezoarticus*. The mitochondrial phylogeny is shown in Figure 3.

**Figure 3.**
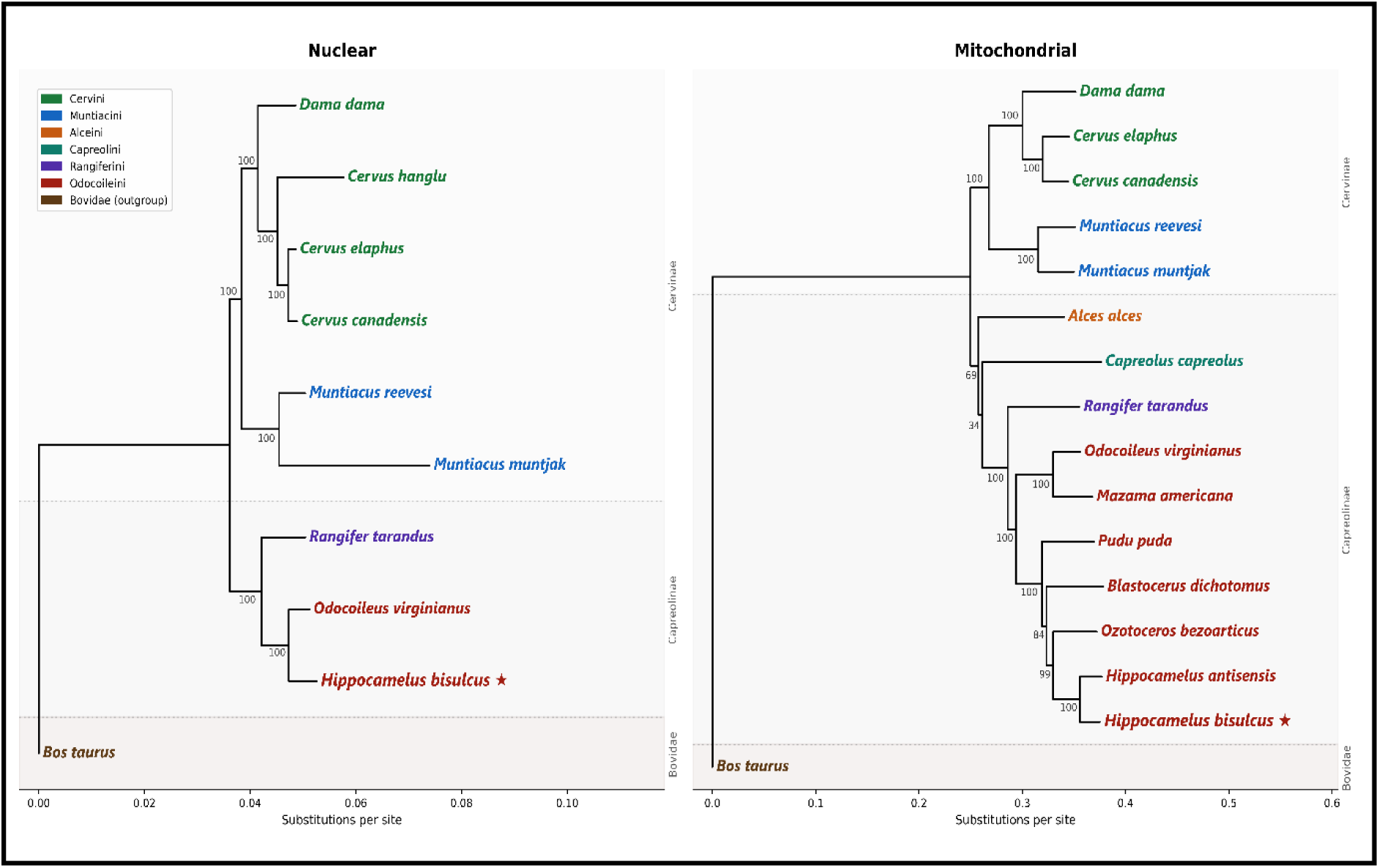
Maximum-likelihood phylogenies of Cervidae based on nuclear and mitochondrial data. The nuclear phylogeny was inferred from concatenated single-copy orthologous proteins across 10 proteomes, whereas the mitochondrial phylogeny was inferred from 16 complete mitochondrial genomes. In both panels, tip labels are colored by tribe, and shaded background bands separated by dotted lines delimit the subfamilies Cervinae and Capreolinae. Bos taurus (Bovidae) served as the outgroup. *Hippocamelus bisulcus* is shown in red and marked with a star (★). Values at internal nodes indicate ultrafast bootstrap support (%), and branch lengths represent substitutions per site.

### Nuclear genome assembly

Given the limited availability of biological samples for this endangered species, together with constraints on sequencing resources, the nuclear genome assembly was generated by combining reads from two individuals, Shehuen providing the primary dataset and Coirón contributing supplementary coverage. To identify the optimal assembly strategy, multiple Flye configurations were benchmarked, varying minimum read length thresholds and overlap parameters, using both the single-individual (Shehuen only) and combined (Shehuen + Coirón) read sets. The best-performing configuration, obtained from the combined read set, yielded a draft nuclear genome of 2.55 Gb distributed across 13,762 contigs, with a contig N50 of 8.71 Mb (L50 = 92), an N90 of 1.23 Mb (L90 = 353), a longest contig of 31.01 Mb, and an overall GC content of 41.68%. BUSCO assessment against the cetartiodactyla_odb10 lineage dataset (n = 13,335 orthologs) indicated 98.8% genome-mode completeness (C:98.8% [S:94.8%, D:4.0%], F:0.3%, M:0.9%). The single-individual assembly (Shehuen only) already achieved high completeness (98.0% BUSCO; S: 94.1%, D: 3.9%) but lower contiguity (contig N50 = 3.13 Mb, longest contig = 11.96 Mb). Although Coirón contributed only ∼15% of the total sequencing data, adding its reads nearly tripled both the contig N50 (3.13 → 8.71 Mb) and the length of the longest contig (11.96 → 31.01 Mb), while completeness increased only marginally (98.0 → 98.8%). The combined assembly showed essentially no change in either total size (2.54 → 2.55 Gb) or duplicated BUSCO content (3.9 → 4.0%) relative to the single-individual assembly, indicating that adding the second individual improved contiguity without evidence of a substantial increase in haplotypic redundancy based on these assembly-wide metrics. Given these results, the combined Shehuen+Coirón assembly was selected as the basis for all downstream analyses.

### Post-assembly processing

#### Duplicate purging and polishing

To remove falsely duplicated sequences arising from unresolved heterozygosity, the Flye assembly was processed with purge_dups, reducing the number of contigs from 13,762 to 10,253 while retaining 2.52 Gb of sequence (N50 = 8.73 Mb, N90 = 1.52 Mb). BUSCO assessment of the purged assembly indicated 98.7% completeness (C:98.7% [S:94.9%, D:3.9%], F:0.4%, M:0.9%), with a slight reduction in duplicated BUSCOs (4.0% to 3.9%) relative to the pre-purge assembly, consistent with the removal of redundant haplotypic sequences. The purged assembly was subsequently polished with Medaka to refine the consensus sequence without altering assembly structure (2.52 Gb across 10,253 contigs; N50 = 8.73 Mb, N90 = 1.52 Mb), and BUSCO completeness remained stable (C:98.7% [S:94.9%, D:3.9%]). Merqury analysis based on the available ONT read k-mers yielded a QV estimate of 47.21 for the raw Flye assembly and 48.11 after Medaka polishing, with an estimated k-mer completeness of 95.63%. Together, these results indicate a modest improvement in consensus accuracy after polishing and high recovery of the sequence content represented in the ONT reads.

### Contamination screening and contig filtering

The polished assembly was screened for foreign sequence contamination using NCBI FCS-GX, which identified six contigs assigned to *Bradyrhizobium* spp., soil-and plant-associated bacteria consistent with incidental environmental exposure during field-based wildlife sampling. These contigs were removed and the assembly was additionally screened for mitochondrial sequences by megablast against the previously assembled *H. bisulcus* mitogenome. The 64 contigs returning hits contained only short, mostly divergent alignment tracts (72–91% identity; median ∼0.6 kb, maximum 11 kb), together with a minority of near-identical tracts (>98% identity, up to 8 kb), all embedded within large nuclear contigs. In every case, the mitochondrial-homologous sequence occurred internally as a NUMT and did not constitute a standalone mitochondrial contig, indicating that the mitogenome was not retained as unintegrated contamination. All contigs shorter than 20 kb were subsequently removed. Despite adaptor trimming during Dorado basecalling, final screening with NCBI FCS-adaptor identified six residual ONT adaptor fragments, totaling 195 bp, which were trimmed using the NCBI clean-genome procedure. The resulting cleaned assembly comprised 1,581 contigs totaling approximately 2.50 Gb, with a contig N50 of 8.75 Mb and a contig N90 of 1.59 Mb. Genome-mode BUSCO completeness remained essentially unchanged at 98.7% (C:98.7% [S:94.9%, D:3.8%], F:0.4%, M:0.9%), indicating that filtering did not measurably affect conserved gene completeness.

### Scaffolding

*Odocoileus virginianus* was used as the scaffolding and annotation reference because it combines phylogenetic proximity to *H. bisulcus* with the availability of a chromosome-level genome assembly and a comprehensive RefSeq annotation, enabling both reference-guided scaffolding and homology-based annotation transfer. Reference-guided scaffolding against this genome organized 94% of the assembly into 36 chromosome-scale pseudomolecules, representing 34 autosomes plus the X and Y chromosomes based on their correspondence with the reference chromosomes. Prior to GenBank submission, the composite RagTag chr0 sequence was decomposed into its constituent unplaced contigs, yielding a final assembly of 342 sequence objects: 36 chromosome-scale pseudomolecules, 119 reference-associated nonchromosomal scaffolds, and 187 unplaced contigs, totaling 2.50 Gb. The final assembly reached a scaffold N50 of 68.56 Mb (L50 = 15), a scaffold N90 of 42.59 Mb (L90 = 33), and a longest scaffold of 111.06 Mb. QUAST comparison against the nuclear *O. virginianus* reference used for scaffolding showed that the final assembly covered 97.4% of the scaffolding reference, with a duplication ratio of 1.005. Genome-mode BUSCO assessment against the cetartiodactyla lineage showed 99.0% completeness (C:99.0% [S:95.2%, D:3.8%], F:0.2%, M:0.7%), compared with 98.7% in the filtered pre-scaffolding assembly.

### Genome annotation

Prior to gene annotation, a *de novo* repeat library was generated with RepeatModeler, identifying 773 repeat families. Subsequent masking with RepeatMasker showed that 38.09% of the final scaffolded assembly consisted of repetitive sequences, dominated by retroelements (35.41%), particularly LINEs (26.41%), including RTE/Bov-B (12.92%) and L1/CIN4 (12.97%), followed by DNA transposons (1.47%) and simple repeats.

Gene annotation was evaluated using two complementary approaches: evidence-guided *ab initio* prediction with BRAKER3 and reference-based annotation transfer with LiftOn. BUSCO completeness was compared using representative proteomes containing the longest protein isoform per gene. BRAKER3, run with *O. virginianus* RNA-seq from two tissues and *O. virginianus* protein sequences as evidence, predicted 18,754 gene models and 23,716 transcripts, with a mean of 9.4 exons per transcript and a mean CDS length of 1.60 kb. Its representative proteome reached 81.8% protein-mode BUSCO completeness (C:81.8% [S:79.0%, D:2.8%], F:1.1%, M:17.0%). In parallel, LiftOn transferred 20,042 protein-coding genes and 54,647 transcripts from the annotated *O. virginianus* reference, with a mean of 13.2 exons per transcript, a mean CDS length of 2.05 kb, and an average of 2.7 transcripts per gene. The representative LiftOn protein set comprised 20,067 sequences and achieved 95.9% BUSCO completeness in protein mode (C:95.9% [S:95.1%, D:0.8%], F:0.7%, M:3.4%). The difference between the 20,067 protein sequences and the 20,042 mRNA-associated gene models reflects 29 models without an extractable CDS and the retention of 54 translated antigen-receptor segments (36 V and 18 C segments). The LiftOn-derived annotation showed substantially greater completeness and fewer missing and duplicated BUSCOs than the BRAKER3 annotation and was therefore selected as the primary gene set for downstream analyses.

### Comparative genomics

Comparative genomic analysis across nine Cervidae proteomes and *Bos taurus* as outgroup yielded 21,979 orthogroups from 228,543 protein sequences. Of these, 13,773 contained representatives from all ten species, and 10,906 consisted entirely of single-copy genes. Of the 20,067 *H. bisulcus* protein sequences, 19,859 (99.0%) were assigned to orthogroups; of these, 19,823 (98.8%) were shared with at least one other species, while 36 (0.2%) formed nine *H. bisulcus*-specific orthogroups; 208 (1.0%) remained unassigned.

The nuclear phylogeny, inferred from the concatenated alignment of single-copy orthologous proteins shared across all species, placed *H. bisulcus* within Odocoileini and recovered *O. virginianus* as its closest sampled relative, with maximal support (ultrafast bootstrap = 100%; Figure 3).

The principal outcomes of the nuclear and mitochondrial genome assembly, curation, and annotation workflows are summarized in Figure 2.

## Discussion

High-quality reference genomes have become increasingly important for conservation genomics, providing the framework required for genome-wide analyses in non-model and threatened species (Supple & Shapiro, 2018; Theißinger et al., 2023). Nevertheless, genomic resources remain sparse and unevenly distributed across Cervidae, limiting the resolution of evolutionary and comparative genomic analyses (Zhong et al., 2026). Here, we present the first nuclear genome assembly for *Hippocamelus bisulcus* and for the genus *Hippocamelus*, together with a complete mitochondrial genome for the species. Despite moderate sequencing depth (∼22.8×), the Oxford Nanopore long-read dataset produced a highly contiguous 2.50-Gb nuclear assembly, with a contig N50 of 8.75 Mb, 99.0% genome-level BUSCO completeness, an estimated k-mer completeness of 95.63%, and an ONT k-mer-based QV estimate of 48.11. Assembly size was comparable to that reported for other cervids, including *Odocoileus virginianus* and *O. hemionus* (Lamb et al., 2021; London et al., 2022). Together with the annotation of 20,042 protein-coding genes and a complete 16,405-bp mitochondrial genome, these results provide a comprehensive genomic foundation for future studies of huemul evolution and conservation.

The quality of the assembly is particularly notable given the constraints associated with sampling an endangered species. The majority of the sequencing data originated from Shehuen, whereas Coirón contributed ∼15% of the combined dataset. Nevertheless, incorporating this additional coverage increased contig N50 from 3.13 to 8.71 Mb and the longest contig from 11.96 to 31.01 Mb, while total assembly size and duplicated BUSCO content remained essentially unchanged. Thus, in this case, the limited contribution of reads from a second conspecific individual substantially improved contiguity without producing the extensive haplotypic redundancy reported for more heterogeneous pooled-sample assemblies (Goldberg et al., 2024). The present assembly, however, should be regarded as a composite, consensus-like reference rather than as a haplotype-resolved diploid genome of either sampled individual. Conventional long-read assemblers commonly collapse homologous haplotypes into a consensus representation, resulting in loss of phase information, whereas robust reconstruction of both haplotypes requires haplotype-aware assembly and sufficient individual-specific sequence and phasing information (Cheng et al., 2021, 2022). A future haplotype-resolved reference for *H. bisulcus* would therefore require sequencing a single individual at sufficient depth using data and phasing strategies specifically designed for diploid assembly.

Reference-guided scaffolding organized 94% of the assembly into 36 chromosome-scale pseudomolecules (34 autosomes plus X and Y), raising the scaffold N50 to 68.56 Mb from a pre-scaffolding contig N50 of 8.75 Mb. These pseudomolecules provide a useful provisional chromosomal framework, but because their assignment and ordering were informed by the *O. virginianus* reference, they remain reference-guided representations rather than independently validated huemul chromosomes, and lineage-specific rearrangements or inversions may go undetected (Udall & Dawe, 2018; Yamaguchi et al., 2021). This caveat is especially relevant in cervids, where physical mapping has revealed misassembled scaffolds even in chromosome-level assemblies (Poisson et al., 2023). Independent long-range evidence, such as Hi-C, optical mapping, linkage, or cytogenetics, will be needed to confirm chromosome organization in *H. bisulcus*. Accordingly, the present resource is best described as a highly complete, reference-guided chromosome-scale assembly.

The close genomic correspondence with *O. virginianus* also facilitated structural annotation. When evaluated using representative proteomes containing the longest protein isoform per gene, LiftOn transferred 20,042 protein-coding genes and recovered 95.9% of the Cetartiodactyla BUSCO set, substantially exceeding the 81.8% obtained with BRAKER3. This difference is consistent with the strengths of homology-based annotation transfer when a well-annotated genome from a closely related species is available, whereas evidence-guided prediction depends strongly on the amount and relevance of the transcriptomic and protein evidence supplied (Chao et al., 2025; Freedman & Sackton, 2025). However, because LiftOn relies on coordinate-based transfer from the *O. virginianus* annotation, it is inherently better suited to recovering conserved gene models than to identifying lineage-specific or substantially diverged genes. Accordingly, the high BUSCO completeness should be interpreted as evidence of strong recovery of the conserved cetartiodactyl gene repertoire rather than as a comprehensive assessment of *H. bisulcus*-specific gene content. Consistently, OrthoFinder assigned 19,823 of the 20,067 *H. bisulcus* protein sequences included in the comparative analysis (98.8%) to orthogroups shared with at least one other species, supporting broad recovery of the conserved cervid protein repertoire. Only 36 proteins (0.2%) occurred in nine *H. bisulcus*-specific orthogroups; given the reference-based origin of the annotation, these should be interpreted cautiously and not as evidence of lineage-specific genes without additional species-specific validation.

The repetitive landscape of the huemul genome was also consistent with that reported for other cervids. Repetitive sequences accounted for 38.09% of the assembly and were dominated by retroelements, particularly LINEs, with RTE/Bov-B and L1/CIN4 representing the major LINE families. Comparable repeat-rich landscapes have been reported across Cervidae, with repetitive content ranging from approximately 36–44% in several deer genomes and LINEs representing a major component of the repetitive fraction (Zhong et al., 2026). These findings place the repetitive content recovered in the huemul assembly within the range reported for other cervid genomes and provide additional evidence that the assembly captures the expected LINE-rich repetitive landscape.

The mitochondrial and nuclear phylogenetic analyses recovered the expected taxonomic affinities of *H. bisulcus*, providing complementary biological validation of the genomic resources generated in this study. The 16,405-bp mitogenome contained the canonical vertebrate complement of 13 protein-coding genes, 22 tRNAs, and two rRNAs (Boore, 1999). A BLASTn comparison identified the *H. antisensis* mitogenome (NC_020711.1) as its closest match, with 96.23% nucleotide identity across the complete sequence. Previous phylogenetic studies based on cytochrome *b* sequences from individual specimens (Duarte et al., 2008; Gutiérrez et al., 2017; González & Barbanti Duarte, 2020), as well as broader analyses incorporating additional mitochondrial markers and a limited number of nuclear loci (Gilbert et al., 2006; Heckeberg, 2020), also placed *H. bisulcus* within the South American odocoileine radiation, although its precise relationships varied among datasets. In the complete-mitogenome phylogeny presented here, the newly assembled *H. bisulcus* mitogenome and the *H. antisensis* NCBI RefSeq mitogenome formed a maximally supported sister pair within a South American odocoileine assemblage that also included *Blastocerus* and *Ozotoceros*. By providing the first complete mitogenome for *H. bisulcus*, our analysis extends previous marker-based evidence and enables comparison between the two recognized *Hippocamelus* species at the complete-mitogenome scale. At the nuclear proteome scale, *H. bisulcus* was likewise recovered within Odocoileini, indicating that the assembled and annotated genome captures the expected phylogenetic signal. Within the available taxonomic sampling, *O. virginianus* was recovered as its closest sampled relative, with maximal support. This relationship should be interpreted in the context of the available taxonomic sampling, because *H. antisensis* and several other South American odocoileines were absent from the nuclear dataset. Although genome assemblies are available for some additional taxa, the absence of comparable annotated proteomes prevented their inclusion in the proteome-based analysis. Genome-scale data and comparable protein annotations from additional Neotropical deer will therefore be required to resolve the nuclear relationships of *Hippocamelus* within the South American odocoileine radiation.

Although future incorporation of higher-depth single-individual sequencing, complementary short-read polishing, independent long-range chromosomal information, and species-specific transcriptomic data would further improve consensus accuracy, haplotype resolution, chromosome validation and gene annotation, the present assembly already provides a high-quality, highly complete and contiguous reference suitable for a broad range of comparative and population-genomic applications. For the huemul, such applications are particularly relevant because the species persists in small and fragmented populations and previous genetic studies have relied primarily on mitochondrial markers and microsatellites (Corti et al., 2011; Smith-Flueck et al., 2025). The genome presented here provides a framework for reference-based population genomic analyses, variant discovery, SNP-panel development, non-invasive genomic monitoring, and future analyses of population structure, effective population size, genomic connectivity, inbreeding, and runs of homozygosity when adequate population-level sequencing becomes available (Supple & Shapiro, 2018; Theißinger et al., 2023). The annotated gene set will additionally enable future investigation of coding variation in pathways of potential conservation relevance, including immune, reproductive, and metabolic functions. Any candidate associations, however, will require validation using appropriate population-level and functional evidence.

More broadly, this study illustrates the potential of Oxford Nanopore long-read sequencing for generating genomic resources under the sampling and logistical constraints that often accompany threatened-species research. Recent studies have demonstrated that ONT long-read sequencing can generate highly contiguous genomic resources for threatened non-model species, including endangered primates and birds, and can be effectively combined with reference-guided scaffolding to generate provisional chromosome-scale organization when appropriate (Hauff et al., 2025; Pozo et al., 2024; Sessi et al., 2026). In the present study, moderate long-read coverage combined with high-accuracy basecalling, polishing, and careful post-assembly curation was sufficient to produce a highly complete and contiguous nuclear genome without complementary short-read sequencing. By moving *H. bisulcus* from marker-based genetic studies to a genome-scale framework, this resource establishes the basis for population and conservation genomics in the species and provides a foundation for incorporating genomic information into the long-term study and management of this endangered Patagonian cervid.

## Data Availability Statement

The nuclear genome assembly and complete mitochondrial genome have been deposited at DDBJ/ENA/GenBank under BioProject PRJNA1509444. The nuclear assembly is registered under WGS accession JCCBWI000000000, and the mitochondrial genome was included as the organellar sequence associated with the same genome submission. The assembly version described in this paper is JCCBWI010000000. Quality-filtered Oxford Nanopore reads are available from the NCBI Sequence Read Archive under accessions SRR40078882 (Shehuen) and SRR40078881 (Coirón), associated with BioSamples SAMN62264853 and SAMN62264854, respectively. Nuclear and mitochondrial annotation files, the custom RepeatModeler library, RepeatMasker outputs, phylogenetic alignments and trees, comparative-genomics results, and associated quality-control files are available from Zenodo at https://doi.org/10.5281/zenodo.22063022. Custom bioinformatic scripts and workflow documentation are available from the project GitHub repository at https://github.com/judeland/huemul-genome-pipeline.

## Ethics Statement

The veterinary and conservation management procedures during which the blood samples were collected were authorized by the Dirección de Fauna y Flora Silvestre de la Provincia del Chubut under Disp. No. 11/2024-DFyFS-MP. No animals were handled or sampled specifically for the present genomic study.

## Acknowledgments

The authors thank Fundación Shoonem, Fundación Temaikèn, and their staff and collaborators for facilitating access to biological samples and for their work in huemul capture, veterinary care, translocation, and management at the Shoonem Breeding and Rehabilitation Center. We acknowledge the Erlenmeyer Stiftung, Switzerland, for supporting the broader huemul conservation initiative, and the DodoBahati Foundation and the Dallas Safari Club Foundation for their financial support of associated logistical activities. We also thank the Wildlife Trust Foundation, Switzerland, for its support with laboratory supplies used in this study. We thank the Dirección de Fauna y Flora Silvestre de la Provincia del Chubut for authorizing and supporting the associated research and conservation activities. We gratefully acknowledge Oxford Nanopore Technologies and New England Biolabs for their support through the ORG.one initiative, including the provision of sequencing consumables and DNA extraction reagents, respectively.

## Author Contributions

Conceptualization, M.J.O.; Methodology, M.J.O.; Software, M.J.O.; Validation, M.J.O.; Formal analysis, M.J.O.; Investigation, M.J.O.; Resources, M.J.O., J.A.M.S.-F., W.T.F., V.P., E.A. and A.V.; Data curation, M.J.O.; Writing—original draft, M.J.O.; Writing—review and editing, M.J.O., J.A.M.S.-F., W.T.F., V.P., E.A. and A.V.; Visualization, M.J.O.; Supervision, M.J.O.; Project administration, M.J.O. and A.V.; Funding acquisition, M.J.O. All authors have read and approved the final version of the manuscript.

## Declaration of generative AI and AI-assisted technologies

During manuscript preparation, the authors used Jenni AI and Elicit to assist with literature discovery and reference organization, and ChatGPT (OpenAI) and Claude (Anthropic) to assist with language editing, manuscript organization, and bioinformatic workflow troubleshooting. All references, analyses, code, and text were independently reviewed and verified by the authors, who take full responsibility for the final content.

